# Expression of a human Gb3/CD77 synthase in insect and human cells: comparison of activity and glycosylation

**DOI:** 10.1101/2025.06.09.658241

**Authors:** Krzysztof Mikolajczyk, Katarzyna Szymczak-Kulus, Anna Bereznicka, Radoslaw Kaczmarek, Lukasz Filip Sobala, Anna Jakubiak-Augustyn, Marcin Czerwinski

**Affiliations:** Laboratory of Glycobiology, Hirszfeld Institute of Immunology and Experimental Therapy, Polish Academy of Sciences, Rudolf Weigl St. 12, 53-114, Wroclaw, Poland; Department of Pediatrics, Indiana University School of Medicine, IUPUI-Wells Center for Pediatric Research, Indianapolis, IN, USA; Department of Lipids and Liposomes, Faculty of Biotechnology, University of Wroclaw Fryderyka Joliot-Curie 14a 50-383 Wroclaw Poland.

**Keywords:** Gb3/CD77 synthase, glycosyltransferase, protein expression, BEVS, HEK293

## Abstract

Glycosylation of proteins can impact their folding, stability, trafficking and enzymatic activity. Human Gb3/CD77 synthase (α1,4-galactosyltransferase, A4galt) has two occupied N-glycosylation sites. Previously, we demonstrated that the activity of recombinant enzyme relies on its N-glycosylation. In this study, we produced soluble recombinant catalytic domain of human Gb3/CD77 synthase in two expression hosts known for different glycosylation patterns: *Trichoplusia ni* insect cells (High Five) and human embryonic kidney cells (Expi293F). The High Five cells generate short oligomannose structures, while the Expi293F cells synthesize complex type glycans. We evaluated the activity of High Five-derived and Expi293F-derived enzymes, characterized the structures of their N-glycans and showed that High Five cells provide a higher amount and activity of the enzyme. Moreover, we used the Expi293F cells to evaluate the N- and C-terminal location of the 6xHis-tag and found that only the N-terminally tagged Expi293F-derived enzyme demonstrated activity. In contrast, the enzyme produced in High Five cells was active despite carrying a C-terminal tag. These findings highlight the role of glycosylation pattern and tag position in the activity of human recombinant glycosyltransferase produced in different hosts.

**Highlights:**

- Active recombinant Gb3/CD77 synthase was obtained in High Five and Expi293F cells.
- High Five cells provided higher recombinant Gb3/CD77 synthase yield than Expi293F.
- Glycosylation pattern of recombinant Gb3/CD77 synthase was host specific.
- Activity of Expi293F-derived enzyme depended on 6xHis-tag localization.

## Introduction

Production of recombinant protein in a heterologous host is increasingly used as a means to obtain functional proteins in high yield. However, the selection of an appropriate expression system is a critical step in the production of recombinant proteins. Expression of recombinant proteins is particularly challenging when the goal is to produce an active enzyme, due to the complexity and diversity of post-translational modifications (PTMs) that can significantly influence protein function. Among these, glycosylation is an essential modification, affecting protein folding, stability, intracellular trafficking and enzymatic activity [1].

Glycosyltransferases (GTs) are a large group of enzymes that transfer carbohydrate moieties to a variety of acceptor molecules, including proteins, lipids and nucleic acids. The products of GT activity play pivotal roles in numerous biological processes, such as cell differentiation, signal transduction, immune response and pathogen infection [1]. Human Gb3/CD77 synthase (α1,4-galactosyltransferase, A4galt, P1/P^k^ synthase; EC 2.4.1.228, UDP-galactose: β-D-galactosyl-β1-R4-α-D-galactosyltransferase), encoded by the *A4GALT* gene, is a unique member of the galactosyltransferase family due to its acceptor promiscuity and variant-dependent specificity. It catalyzes the synthesis of Galα1→4Gal structures on glycosphingolipids (GSLs) and N-glycans, leading to the synthesis of Ga2 (galabiosylceramide), Gb3 (globotriaosylceramide, CD77, P^k^ antigen) and P1 antigen [2–4]. However, a variant form of Gb3/CD77 synthase, harboring a p.Q211E substitution, demonstrates a broadened acceptor specificity and can also synthesize the unusual Galα1→4GalNAc moieties of GSL-based NOR antigens. These glycan structures represent histo-blood group antigens belonging to the P1PK group system (International Society of Blood Transfusion, System no. 003) [4,5]. Furthermore, the products of human Gb3/CD77 synthase serve as receptors for pathogens and toxins, such as Shiga toxins (Stxs) released by enterohemorrhagic *Escherichia coli* and *Shigella dysenteriae* serotype 1 [4,5]. Moreover, accumulation of Gb3 (and of Ga2 and the P1 antigen, to a lesser extent) underlies Anderson-Fabry disease (OMIM #301500) caused by α-galactosidase A deficiency [6]. Elevated biosynthesis of Gb3 is also associated with the progression of colorectal, gastric, pancreatic, and ovarian cancer, indicating its role in carcinogenesis [4].

Human Gb3/CD77 synthase is a type II transmembrane protein with a C-terminal catalytic domain facing the Golgi lumen [4]. The polypeptide chain of Gb3/CD77 synthase consists of 353 aa and contains two occupied N-glycosylation sites at positions N_121_ and N_203_. Our previous studies demonstrated that N-glycosylation is essential for Gb3/CD77 synthase activity, subcellular localization and solubility [7]. In particular, the N-glycan at position N_203_ affects enzyme activity, solubility and its Golgi localization, while the glycan at position N_121_ appears to modulate enzymatic activity [7,8].

In this study, we attempted to compare two expression systems: insect cells *Trichoplusia ni* (High Five) and human embryonic kidney cells (Expi293F), for producing the catalytic domain of human Gb3/CD77 synthase (the wild-type enzyme is a type II membrane protein). To enhance its solubility, we employed a truncated form of the enzyme, encompassing amino acids 44-353, corresponding to the stem and the catalytic domain, but excluding the N-terminal cytoplasmic tail and the highly hydrophobic transmembrane domain. Both High Five and Expi293F cells contain the post-translational modification capabilities for the synthesis of glycosylated human proteins, thereby increasing the likelihood of producing a correctly folded and functional enzyme [9]. We determined the activity of recombinant Gb3/CD77 synthases and investigated their N-glycan structures using lectin blotting and MALDI-TOF mass spectrometry. Additionally, we assessed how the position of the 6xHis-tag at the N- and C-terminus affects purification efficiency and enzymatic activity of the recombinant protein produced in Expi293F cells. The successful production of a soluble and active form of Gb3/CD77 synthase in high yield is a critical step toward resolving its three-dimensional structure and gaining deeper insights into the functional properties of this enzyme.

## Materials and methods

### Cell lines, E. coli strains and expression vectors

The following cell lines were used for recombinant human Gb3/CD77 synthase overexpression: *Trichoplusia ni* High Five cells (BTI-TN-5B1-4) (American Type Culture Collection, ATCC, Rockville, MD, USA) and human embryonic kidney 293 (HEK293)-derived Expi293F™ cells (Thermo Fisher Scientific, Waltham, MA, USA). Vectors encoding recombinant human Gb3/CD77 synthase were amplified in *E. coli* XL-1 Blue (Stratagene, La Jolla, CA, USA) for use in High Five cells. For expression in the human Expi293F cell system, ElectroMAX™ DH5α-E Competent Cells (Thermo Fisher Scientific, Waltham, MA, USA) were employed. Vector pGEM-T Easy (Promega, Madison, WI) containing the *A4GALT* gene construct was used to produce the recombinant baculovirus.

### Amplification of A4GALT ORF

A fragment of the *A4GALT* gene (GenBank nucleotide sequence databases with accession number NG_007495.2, NCBI, https://www.ncbi.nlm.nih.gov/) spanning nucleotides 130-1059 of the open reading frame (ORF) was cloned into all expression vectors used in this study. The *A4GALT* ORF fragment encoding a catalytic domain (amino acids from 44 to 353) of the consensus (contains c.631C) enzyme, devoid of transmembrane and cytosol domains, was selected to obtain a soluble protein. All expression vectors used in this study were purified using the Plasmid Maxi Kit (Qiagen, Venlo, the Netherlands) according to the manufacturer’s instruction. The *A4GALT* ORF fragment was amplified by PCR performed in an MJ Mini gradient PCR thermal cycler (BioRad, Hercules, CA, USA). About 20 μl of reaction mixture contained approx. 200 ng of the template DNA (pCAG vector containing the full-length *A4GALT* ORF) [5], 0.2 mM forward and reverse primers, 0.2 mM dNTPs, 1.5 mM MgCl_2_, HF polymerase buffer (1:5 dilution), 1 unit of Phusion High-Fidelity DNA Polymerase (Thermo Fisher Scientific, Waltham, MA, USA). The primers introduced restriction endonuclease sites as well as N- or C-terminal tags in frame with *A4GALT* ORF (primer sequences listed in Table S1). The amplified DNA fragments were purified with a gel extraction kit (Gel-Out, A&A Biotechnology, Gdynia, Poland) and their oligonucleotide sequences were verified by sequencing (Genomed, Warsaw, Poland).

### Expression of the recombinant catalytic domain of human Gb3/CD77 synthase in High Five cells

An insect codon-optimized fragment of *A4GALT* ORF encoding the catalytic domain of human Gb3-CD77 synthase (encompassing amino acids 44-353), a C-terminal c-myc tag, and a 6xHis-tag was cloned into the pGEM-T Easy vector. The TA cloning procedure was performed according to the manufacturer’s protocol (Promega, Madison, WI). A resulting vector containing the *A4GALT* ORF fragment was used to obtain high-titer suspensions of recombinant baculovirus ordered from GenScript (Piscataway, NJ), which harbored the cloned *A4GALT*-c-myc-6His fragment downstream of the gp67 secretion signal.

Suspension cultures of High Five cells (BTI-TN-5B1-4) were maintained in serum-free medium ESF 921 medium (Expression Systems, Davis, CA, USA) supplemented with penicillium/streptomycin/amphotericin B (100 X solution, Thermo Fisher Scientific, Waltham, MA, USA), at 27 °C as static (in ventilated tissue culture flasks) or agitated (110 rpm, in Erlenmeyer flasks) cultures. The suspension cultures of High Five cells (3-4 x 10^6^/ml) were infected with the recombinant baculovirus at a multiplicity of infection (MOI) of 5 virus particles per High Five cell. 48 h after infection, the culture was centrifuged (15 min, 7500 x g, 4 °C; centrifuge J2-MC, Beckman Coulter) and the supernatant was collected. After concentration, the supernatant was used for the purification of recombinant Gb3/CD77 synthase.

### Expression of the recombinant catalytic domain of human Gb3/CD77 synthase in Expi293F™ cells

A fragment of human *A4GALT* ORF encoding the catalytic domain of human Gb3-CD77 synthase (encompassing amino acids 44-353), a signal peptide and a 6xHis-tag was cloned into the EBA181-Bio vector backbone (Addgene plasmid #47744; https://www.addgene.org/47744/; RRID: Addgene_47744) [10], which is an expression plasmid for transfection of Expi293F™ cells. The *A4GALT* ORF fragment was cloned into XhoI-NotI restriction sites using primers A4GHEKNTAGsense and A4GHEKNTAGanti for expression of protein with N-terminal localized tags, and A4GHEKCTAGsense and A4GHEKCTAGanti for production of C-terminally tagged protein (primer sequences listed in Table S1).

The Expi293F™ cells were grown in Expi293F medium (Thermo Fisher Scientific, Waltham, MA, USA) at 37 °C, 125 rpm, 8% CO_2_ atmosphere and 80% humidity in Erlenmeyer flasks with ventilated caps (Corning® Erlenmeyer sterile polycarbonate with 0.2 μm ventilated caps) in CO_2_ incubator (PHBI) with orbital shaker (Celltron, Infors HT). The cell culture at a density of 2 x 10^6^ cells/ml was transfected with preformed DNA:Expifectamine (ratio: 1:3) complexes at 1 µg/ml DNA (final concentration) in Opti-MEM™ culture medium (Thermo Fisher Scientific, Waltham, MA, USA). After adding expression enhancers 1 and 2, the culturing temperature was lowered to 32 °C, and incubation was continued until cell viability decreased to 70%.

### Purification of the recombinant catalytic domain of human Gb3/CD77 synthase

Duration of protein production varied between expression systems; for High Five cells, 48 h and ∼120 h for Expi293F cells. The culture supernatant was collected and purified using HisPur™ agarose resin (Thermo Fisher Scientific, Waltham, MA, USA). Culture supernatant was filtered through Stericup® Filter Units with 0.22 µm cut-off (Merck), dialyzed against Ni-NTA purification buffer (50 mM NaH_2_PO_4_, 500 mM NaCl, pH 8.0) for two weeks and loaded into the HisPur^TM^ Resin column. The column was washed with 500-1000 mL of Ni-NTA purification buffer. The bound protein was eluted stepwise by increasing imidazole concentration (20 mM, 50 mM, 100 mM, 125 mM, 150 mM, 175 mM, 200 mM and 500 mM) in the Ni-NTA purification buffer. The fractions eluted with 100 and 200 mM imidazole were pooled, dialyzed against TBS (50 mM Tris-HCl, 150 mM NaCl, pH 7.4), concentrated 20 times using centrifugal filter units (Amicon Ultra 10000 molecular weight cut-off, EMD Millipore, Billerica, MA) and supplemented with 50% glycerol for storage in −80 °C. Protein concentrations were determined using a Nanodrop (Thermo Fisher Scientific, Waltham, MA, USA).

### SDS-PAGE and western blotting

The recombinant proteins were separated in the presence of SDS (Roth) using a 10% polyacrylamide gel and visualized with Coomassie Brilliant Blue R-250 (Roth, Karlsruhe, Germany) or transferred to the nitrocellulose membrane (Roth, Karlsruhe, Germany) as described [7]. The PageRuler Prestained Protein Ladder (Thermo Fisher Scientific, Waltham, MA, USA) was used as a protein standard. The proteins fractionated by SDS-PAGE were transferred to the nitrocellulose membrane and detected with mouse anti-c-myc (hybridoma supernatant diluted 1:10, clone 9E10, obtained from ATCC) or anti-6x-His antibody (clone HIS.H8, Thermo Fisher Scientific, Waltham, MA, USA). Goat anti-mouse IgG (H + L) conjugated with alkaline phosphatase was used as a secondary antibody (Thermo Fisher Scientific, Waltham, MA, USA). Hybridoma supernatant and all antibodies were diluted in TBS + 1% BSA (m/v).

### Enzymatic activity evaluation

The enzymatic activity was evaluated by ELISA with oligosaccharide-polyacrylamide (PAA) conjugates as described before [11]. UDP-Gal (Sigma-Aldrich, St. Louis, MO) was used as a donor, and Galβ1→4Glc-PAA (Lac-PAA, precursor to Gb3) was used as an acceptor. ELISA microtiter plates (Nunc, MaxiSorp, Roskilde, Denmark) were coated overnight at 4 °C with conjugates (2 μg/well) in carbonate/bicarbonate buffer (50 mM Na_2_CO_3_, 50 mM NaHCO_3_, pH 9.6). Enzyme samples (2 μg/well) in cacodylate buffer containing Mn^2+^ cations and the donor substrate (50 mM sodium cacodylate, 14 mM MnCl_2_, 200 μM UDP-Gal, pH 6.32) were loaded in triplicate. The reactions were run for 100 min at 37 °C. The plates were then washed twice with distilled water and thrice with PBS-T (137 mM NaCl, 2.7 mM KCl, 10 MM Na_2_HPO_4_, 1.8 mM KH_2_PO_4_, 0.05% Tween 20) and blocked with 5% BSA in PBS-T. Next, mouse anti-P1 (clone 650, Ce-Immundiagnostika (Eschelbronn, Germany) antibody (diluted 1:100) which recognizes the reaction products was added and incubated for 90 min in room temperature, followed by sequential 1 h incubation with biotinylated anti-mouse or anti-human Ig antibody (each diluted 1:1000, Thermo Fisher Scientific, Waltham, MA, USA) and ExtrAvidin-alkaline phosphatase conjugate (diluted 1:10000, Pierce Rockford, IL, USA). The wash steps were carried out using PBS-T + 1% BSA (m/v). Finally, color reactions were developed with *p*-nitrophenyl phosphate (1 mg/ml in carbonate/bicarbonate buffer with 1 mM MgCl_2_) (Sigma-Aldrich, St. Louis, MO). Plates were read using a 2300 EnSpire Multilabel Reader (PerkinElmer, Waltham, MA) at 405 nm at several time points within 1 h. Data were analyzed using Microsoft Office Excel (Microsoft Corp, Redmond, WA). Negative controls were set up by adding incomplete reaction mixtures (lacking UDP-Gal or enzyme) to coated wells, by adding complete reaction mixtures to uncoated wells or by omitting primary or secondary antibodies.

### Analysis of N-glycans from the recombinant catalytic domain of human Gb3/CD77 synthase

Lectin blotting was performed as previously described in [12]. Each line contained 20 μg of untreated or deglycosylated recombinant catalytic domain of human Gb3/CD77 synthase, which was subjected to SDS-PAGE on a 10% Tris-glycine gel. After electrophoretic transfer, the nitrocellulose membrane was blocked in 5% BSA in TBS overnight at 4 °C. The membrane was washed with TBS and then incubated in a 5 μg/ml solution of biotinylated lectin in the appropriate buffer (used lectins and buffers listed in Table S2) for 1 h. The membrane was subsequently washed and incubated for 1 h with ExtrAvidin conjugated with alkaline phosphatase (Pierce, Rockford, IL, USA), which was diluted 1:5000 in TBS + 1% BSA (m/v) + 0.05% Tween 20 (v/v). The protein bands were visualized using the BCIP/NBT reaction.

N-glycans from the recombinant human Gb3/CD77 synthase obtained from High Five and Expi293F cells were released with PNGase F treatment. 50 µg of the glycoprotein was mixed 1:1 with denaturation buffer (2% SDS m/v, 2% β-mercaptoethanol v/v in PBS, pH 7.4). After incubation at 98 °C for 20 min, the samples were centrifuged at 5,000 g, 1 min. The supernatant was mixed with 5 μl of 10% NP-40, 2.5 μl (25 U) of PNGase F (Promega) and PBS, pH 7.4, up to a final volume of 50 μl. PNGase F treatment was performed overnight at 37 °C. The released N-glycans were purified with the use of Sep-Pak C18 cartridges (Waters), pre-equilibrated in methanol (10 column volumes, CV) and 0.5% acetic acid (5 CV). N-glycans were eluted with 0.5% acetic acid and lyophilized. For MALDI-TOF analysis, 0.8 µl of N-glycans dissolved in ultrapure water was mixed 1:1 on the MALDI target plate (MTP 384, Bruker) with the reactive matrix - anthranilic acid (2-AA, A89855, Sigma-Aldrich, St. Louis, MO). The matrix was dissolved to 10 mg/ml in 900:100:2.5 v/v methanol:water:formic acid. MALDI-TOF analysis was performed using a Bruker UltrafleXtreme instrument in negative reflectron mode. The target spot was shot in multiple places to average the effect of random local concentration variations of co-crystallized oligosaccharides on peak intensities.

## Results

### Expression and characterization of the recombinant catalytic domain of human Gb3/CD77 synthase in High Five cells

For the production of human Gb3/CD77 synthase catalytic domain in insect cells, we utilized the baculovirus expression vector system (BEVS) with High Five (*T. ni*) cells as a host. The recombinant baculovirus encoded a truncated form of human Gb3/CD77 synthase containing the catalytic domain of the enzyme, followed by a C-terminal c-myc and a 6xHis-tag. A schematic summary of enzyme expression in the BEVS approach is shown in Fig. 1.

**Figure 1.**
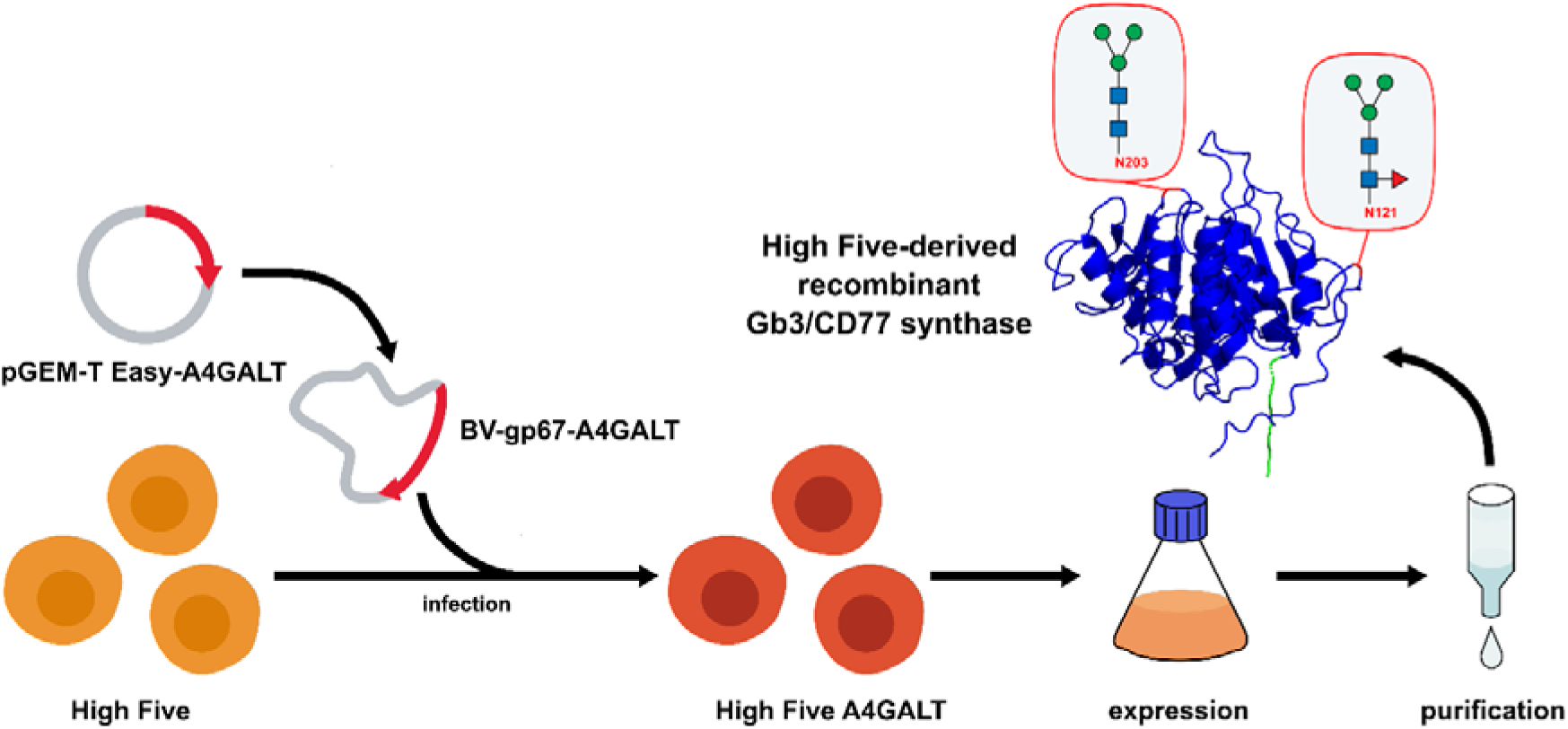
Schematic summary of recombinant catalytic domain of human Gb3/CD77 synthase expression in High Five cells employing the Baculovirus Expression Vector System (BEVS).

Protein purification was performed using HisPur™ agarose resin with stringent washing steps (10 mM or 20 mM imidazole) to reduce non-specific protein binding. Recombinant Gb3/CD77 synthase was eluted from the column with a stepwise rising imidazole concentration (from 20 mM to 500 mM) in Ni-NTA elution buffer. CBB staining exhibited that fractions eluted with 20 mM and 50 mM imidazole contained the highest overall protein levels, including contamination proteins (Fig. 2A). However, fractions eluted with 100-200 mM imidazole yielded higher purity of the recombinant proteins, albeit at lower protein concentrations (Fig. 2A-B). Western blotting analysis using an anti-c-myc antibody (clone 9E10) revealed two forms of the recombinant Gb3/CD77 synthase, a predominant band of approx. 40 kDa corresponding to the monomeric form of the enzyme, and a band of approx. 80 kDa, likely representing a homodimer (Fig. 2B-C).

**Figure 2.**
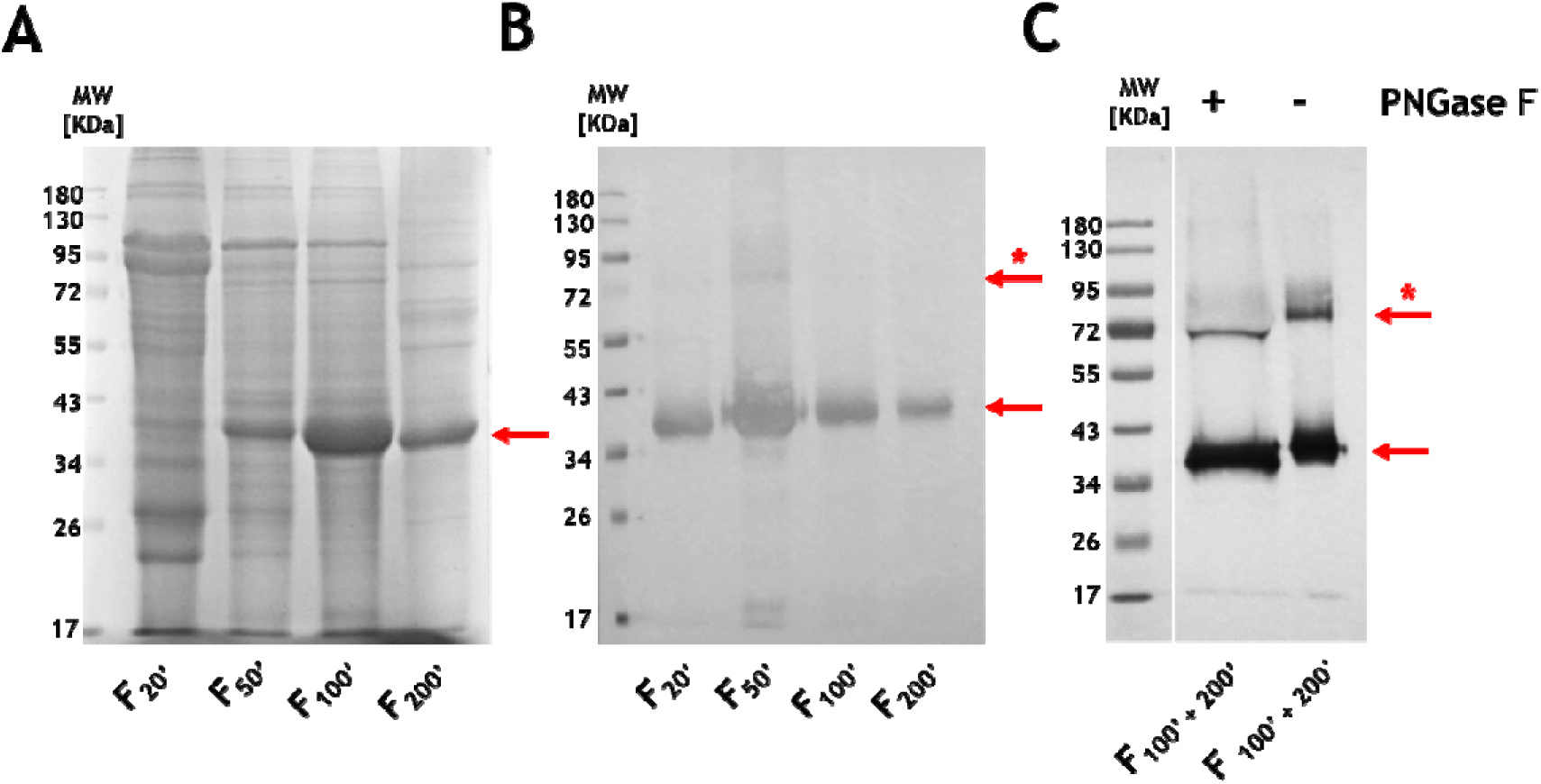
Identification and purity determination for the recombinant catalytic domain of human Gb3/CD77 synthase expressed in the Baculovirus Expression Vector System (BEVS) system using High Five (*T. ni*) cells. Coomassie Brilliant Blue (CBB) staining (**A**) and immunoblotting (**B**) with anti-c-myc antibody (clone 9E10) of the recombinant catalytic domain alongside a molecular weight marker. The presence of N-glycans in recombinant enzyme was examined by treatment of 100’ + 200’ elution fraction with PNGase F and evaluation of the enzyme molecular weight between treated (+) and non-treated (-) samples using western blotting (with anti-c-myc antibody) (C). 20’, 50’, 100’, 200’, elution fraction with 20, 50, 100 and 200 mM imidazole, respectively. The red arrow indicates bands corresponding to the monomer and the red arrow with an asterisk points to the dimeric form of the enzyme.

To evaluate the presence of N-glycan chains in the recombinant catalytic domain of human Gb3/CD77 synthase, the enzyme fraction eluted with 100-200 mM imidazole was treated with PNGase F, which cleaves off N-linked glycans. Western blotting analysis of the treated samples showed a shift in molecular weight from approx. 40 kDa (glycosylated form) to 36 kDa (deglycosylated form), indicating that Gb3/CD77 synthase produced in insect cells is N-glycosylated (Fig. 2C).

Enzymatic activity of the same pooled fractions (100-200 mM imidazole) was assessed using Lac-PAA (Gb3 precursor) as the acceptor substrate. Reaction product formation was detected with the anti-P1 monoclonal antibody (clone 650), which specifically recognizes the Gb3 trisaccharide. The tested enzyme fractions demonstrated Gb3 synthase activity (Fig. 5A, Table 1).

**Table 1.** The comparison of different protein expression systems used for obtaining the recombinant catalytic domain of human Gb3/CD77 synthase.

| Feature | High Five | Expi293F |
| --- | --- | --- |
| **Cells origin** | Insect | Human |
| **Culture type** | Suspension |  |
| **Transfection method** | Recombinant baculovirus | Chemical |
| **Purification method** | Affinity chromatography (Ni<sup>2+</sup>-NTA resin) |  |
| **Protein yield [mg/L]<sup>a</sup>** | 3.5 | 0.5 |
| **N-glycosylation** | Yes (paucimannose-type) | Yes (complex type) |
| **Activity** | Yes | Yes (N-tagged),<br>no (C-tagged) |
| **Cost** | ++ | +++<sup>b</sup> |
| **Effort required** | + (+++<sup>c</sup>) | + |

<sup>a</sup> an estimated enzyme efficiency production from 1 L of culture supernatant (it included contamination with other proteins purified and recombinant human Gb3/CD77 synthase).
<sup>b</sup> the cost of protein production included a commercially available kit that consists of the culture and transfection media and transfection reagents.
<sup>c</sup> when the recombinant baculovirus is made in-house and its titer is determined.

### Expression and characterization of the recombinant catalytic domain of human Gb3/CD77 synthase in Expi293F cells

To produce the recombinant human Gb3/CD77 synthase in the Expi293F cells, constructs encoding the enzyme catalytic domain and a 6xHis-tag were generated. To assess the influence of tag position on enzyme activity and purification efficiency, the tags were added to either the N- or C-terminus of the expression construct. A schematic summary of enzyme expression in Expi293F cells is presented in Fig. 3.

**Figure 3.**
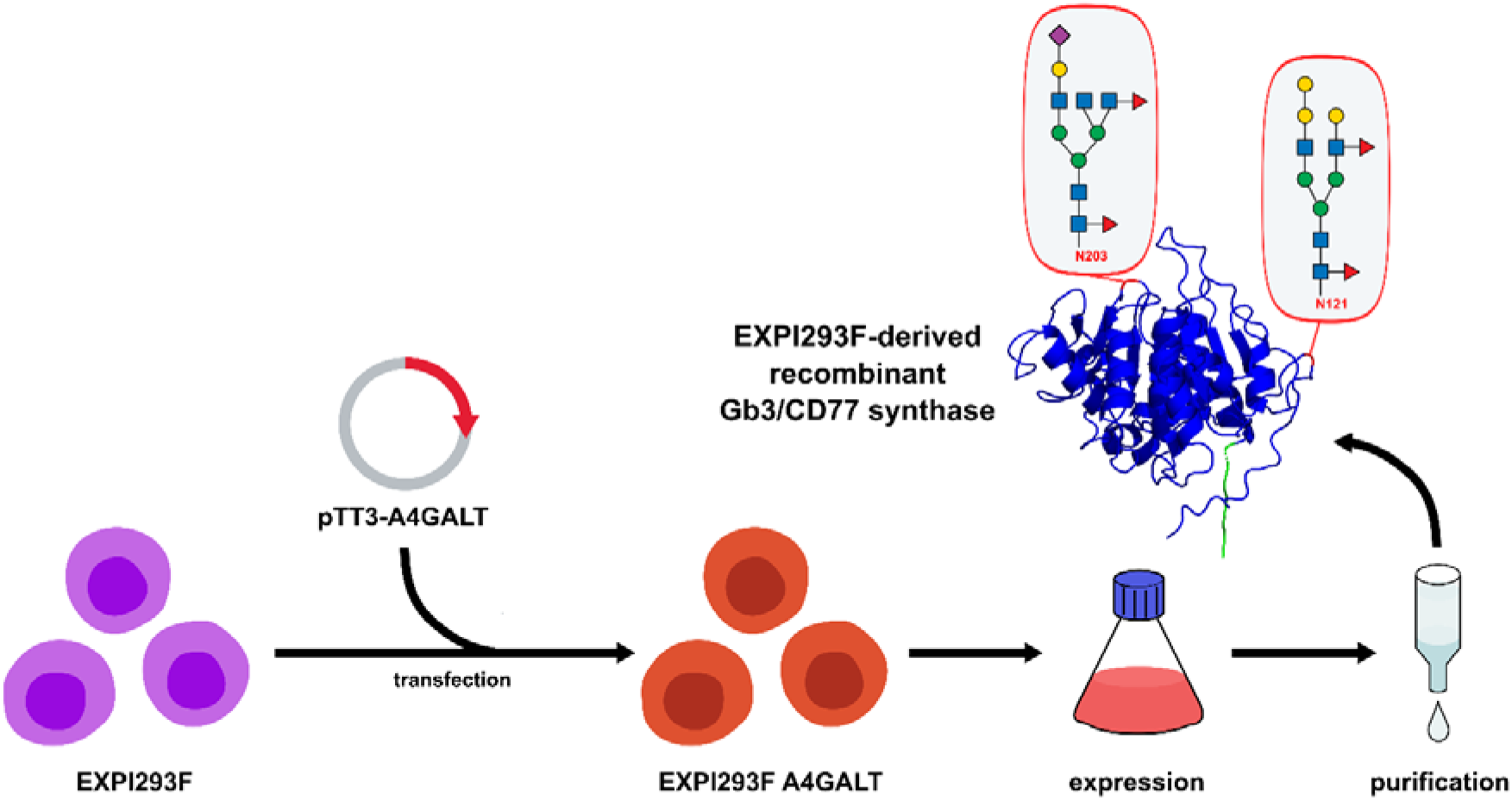
Schematic workflow summary of expression of the recombinant catalytic domain of human Gb3/CD77 synthase in Expi293F cells.

Protein purification was carried out using HisPur^TM^ agarose resin pre-equilibrated by extensive washing with 10 mM imidazole to minimize co-purifying contaminants. The recombinant catalytic domain of human Gb3/CD77 synthase was eluted with stepwise increasing imidazole concentration (20 mM to 500 mM) in Ni-NTA elution buffer. For both C- and N-tagged proteins, SDS-PAGE and CBB staining revealed that fractions eluted with 100-200 mM imidazole exhibited the highest purity, while fractions eluted with 20-50 mM contained significant levels of contaminant proteins (C-tagged enzyme: Fig. 4A, N-tagged enzyme: Fig. 4D). Western blotting analysis using an anti-6xHis-tag antibody confirmed the presence of recombinant human Gb3/CD77 synthase in all analyzed protein fractions (C-tagged enzyme: Fig. 4B, N-tagged enzyme: Fig. 4E).

**Figure 4.**
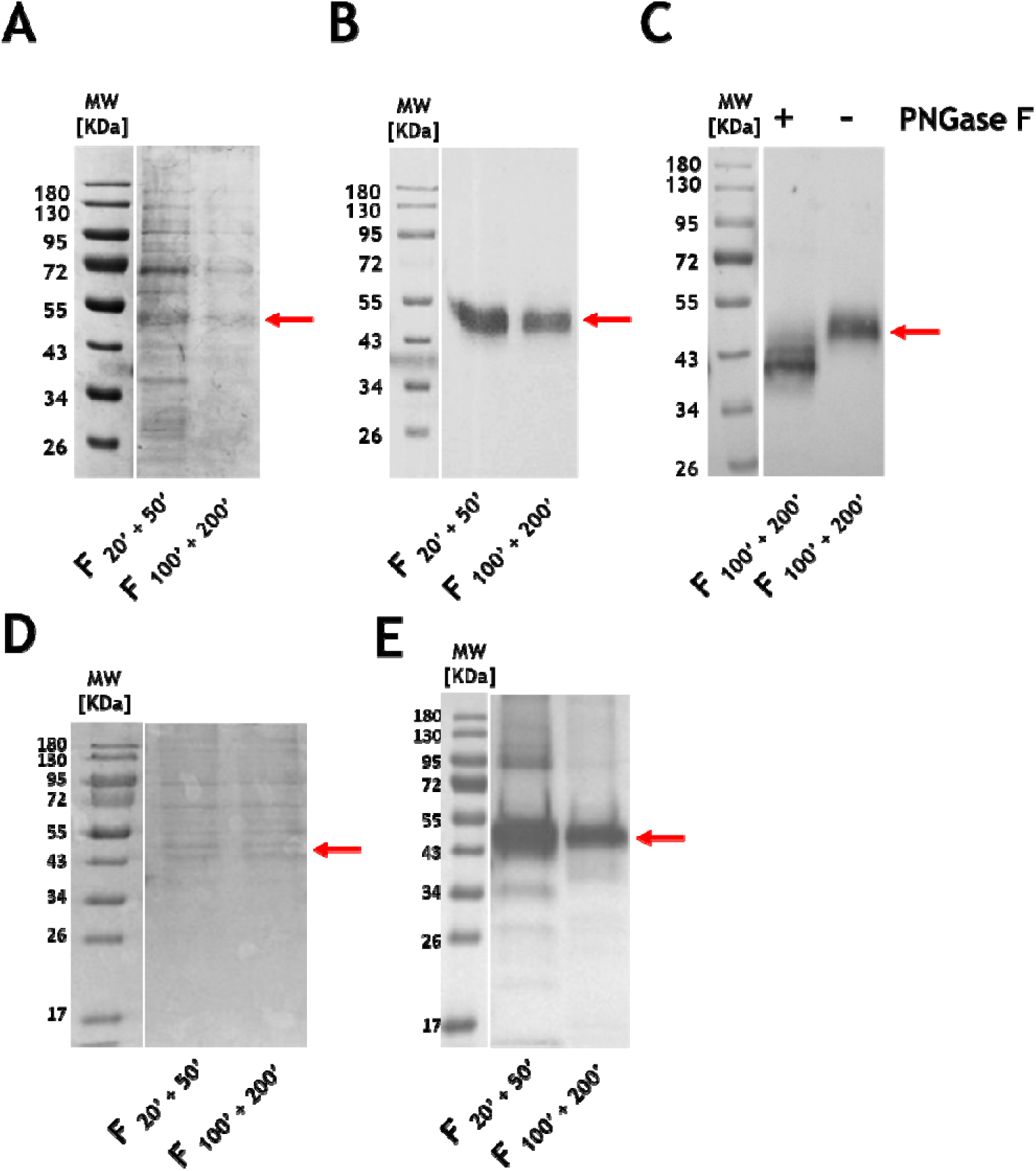
Identification and purity determination for recombinant catalytic domain of human Gb3/CD77 synthase with His-Tag at C- (**A** and **B**) or N-terminus (**D** and **E**), was expressed using human embryonic kidney cells (Expi293F cells). Coomassie Brilliant Blue (CBB) staining (**A** and **D**) and immunoblotting (**B** and **E**) with anti-6x-His antibody (HIS.H8 clone) of recombinant catalytic domain alongside a molecular weight marker are shown. The presence of N-glycans in recombinant enzyme was examined by treatment of 100’ + 200’ elution fraction of C-tagged recombinant catalytic domain of Gb3/CD77 synthase with PNGase F and evaluation of the enzyme molecular weight between treated (+) and non-treated (-) samples using western blotting (with anti-6x-His antibody, clone HIS.H8) (**C**). 20’ + 50’, combined elution fractions with 20 mM and 50 mM imidazole, respectively; 100’ + 200’, combined elution fractions with 100 mM and 200 mM imidazole, respectively. The red arrow indicates bands corresponding to the monomer.

PNGase F digestion indicated the existence of glycosylated (50 kDa) and deglycosylated (43 kDa) forms of recombinant human Gb3/CD77 synthase (Fig. 4C). An additional band at 45 kDa was also observed, suggesting the presence of partially deglycosylated enzyme form (Fig. 4C). Enzymatic assays using Lac-PAA as an acceptor substrate demonstrated activity only for the N-terminally tagged recombinant catalytic domain, with no detectable activity in the C-terminally tagged variant (Fig. 5B, Table 1), likely due to steric interference of the C-terminal tag with the catalytic center. In summary, expression in human cells enabled the production of a soluble, catalytically active form of human Gb3/CD77 synthase (N-terminally tagged).

**Figure 5.**
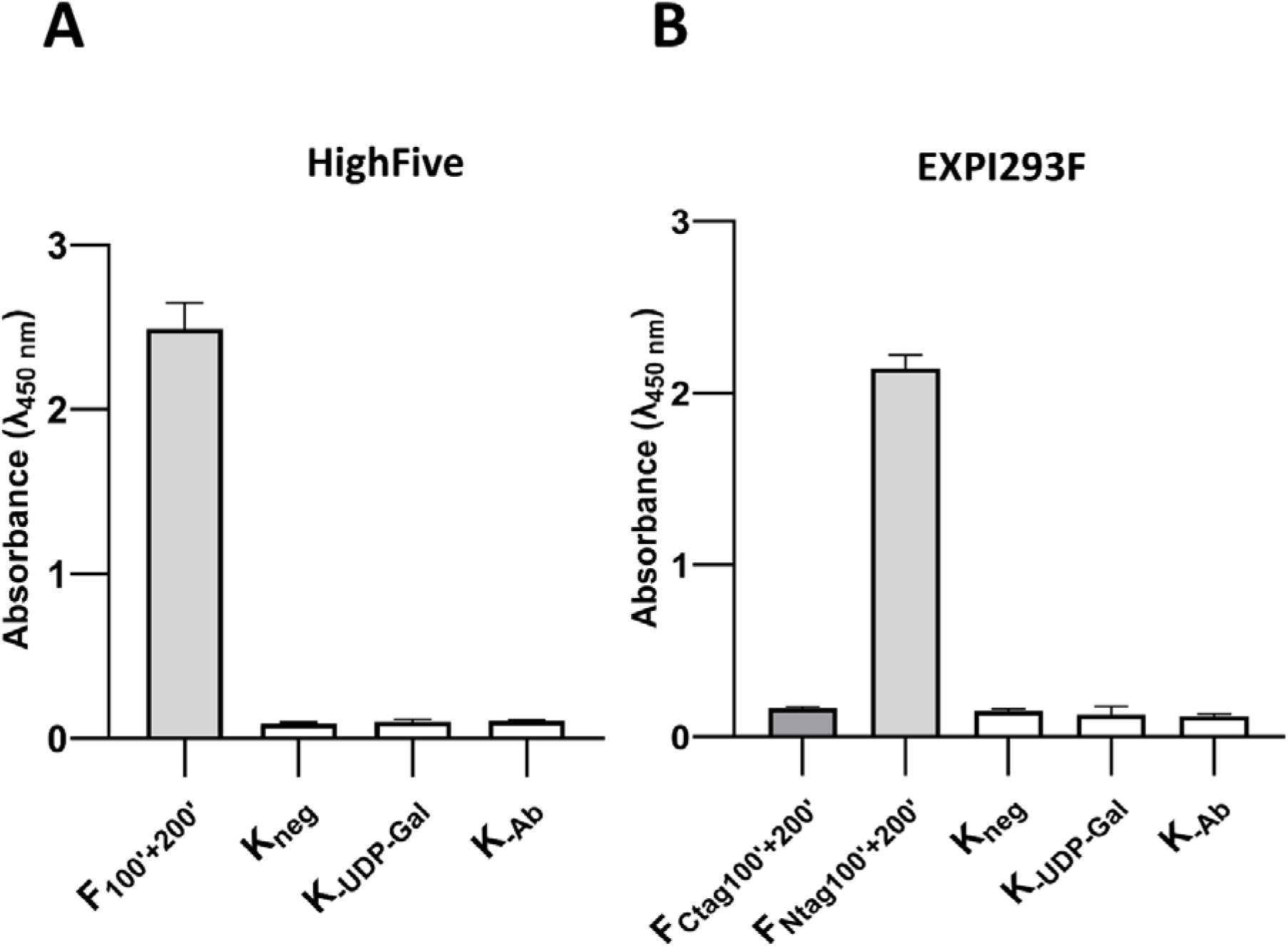
Evaluation of the enzymatic activity of the purified recombinant catalytic domain of Gb3/CD77 synthase obtained using High Five (**A**) and Expi293F (**B**) cells by ELISA. Activities of the recombinant enzymes were evaluated toward Lac-PAA conjugate (a precursor of Gb3) used as the acceptor. F_100’ + 200’_ combined elution fractions with 100 mM and 200 mM imidazole, respectively; K_Neg_, no enzyme control; K_-UDP-Gal_, no donor control; K_-Ab_, no antibody control.

### Analysis of N-glycan structures in the recombinant catalytic domain of human Gb3/CD77 synthase

Lectin blotting and MALDI-TOF mass spectrometry analysis of released N-glycans were applied to characterize the glycan structures in the recombinant enzymes produced in insect and human cells. Lectin blots revealed differences in glycosylation patterns between the two expression systems. The bands corresponding to the High Five-derived enzyme were predominantly stained by Con A (specific for oligomannose- and hybrid-type glycans), LCA (which recognizes oligomannose and core fucosylated N-glycans), and RCA I (which binds to terminal galactose residues) (Fig. 6A) [13]. These findings suggest that the insect cell-derived enzyme carries a mixture of high-mannose and fucosylated structures, with terminal galactosylation. Only a weak band was observed in SNA staining, which binds α2,6-linked sialic acid, indicating the low level or lack of sialylation in the High Five-produced enzyme (Fig. 6A).

**Figure 6.**
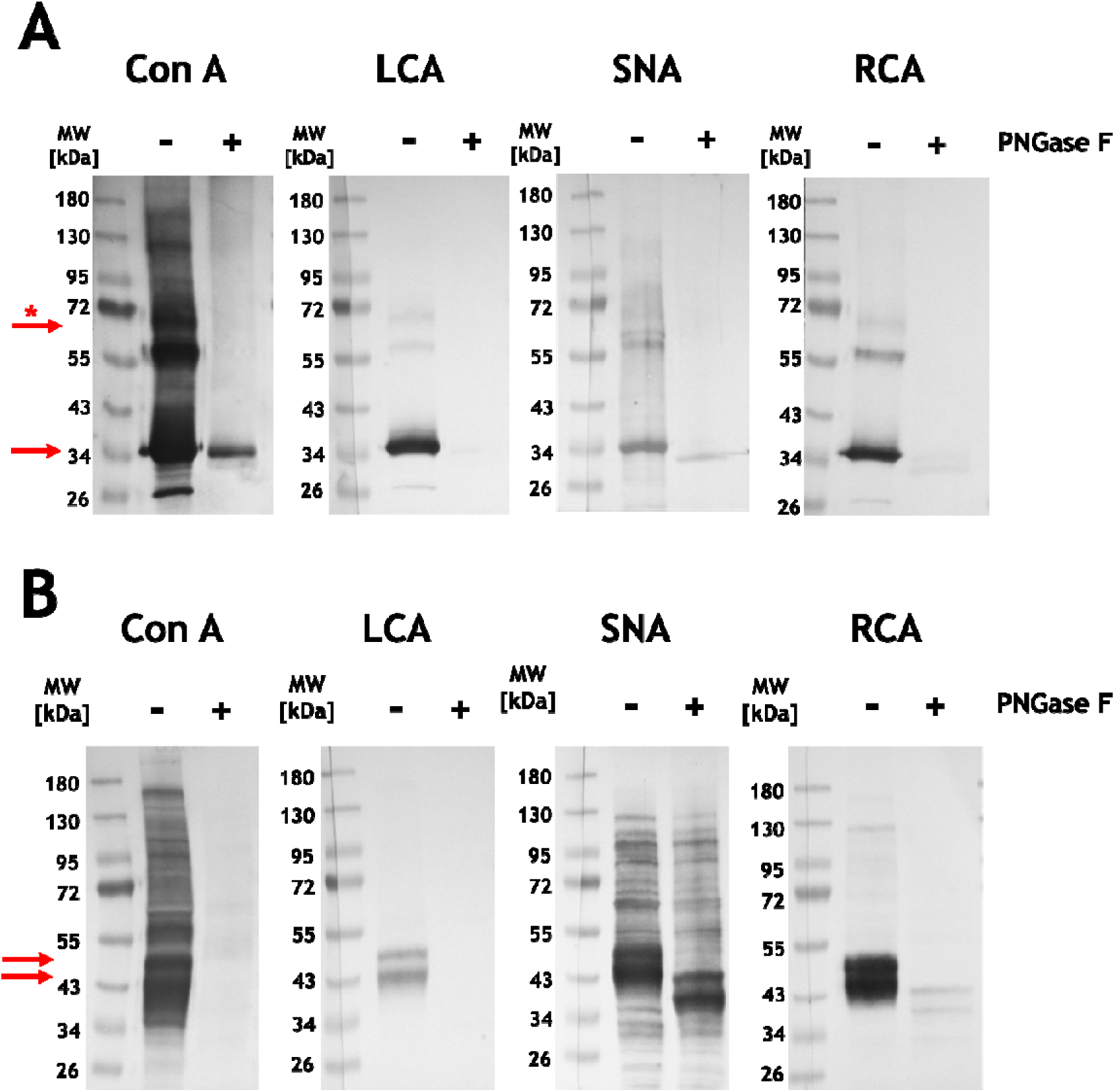
Lectin blot analysis of recombinant catalytic domain of human Gb3/CD77 synthase (100’ protein fraction eluted using 100 mM imidazole) obtained using High Five (*T. ni*) cells (**A**) and human embryonic kidney cells (**B**, Expi293F, 100’ + 200’ protein fractions eluted using 100 mM and 200 mM imidazole), stained with Con A, LCA, SNA and RCA I (lectins specificity and buffers listed in Table S2). The analyzed recombinant proteins were treated (+) or non-treated (-) with PNGase F. The presence of the protein bands after PNGase F treatment indicated partial deglycosylation. The red arrow indicates bands corresponding to the monomer and the red arrow with an asterisk points to the dimeric form of the enzyme.

In contrast, Gb3/CD77 synthase produced in Expi293F cells was strongly bound by Con A, SNA, and RCA I, with relatively weak binding to LCA (Fig. 6B). The pronounced SNA binding indicates the presence of sialylated N-glycans, consistent with glycosylation profiles typically associated with mammalian expression systems. The reduced interaction with LCA suggests a lower abundance of core fucosylation, or a prevalence of glycan structures with lower affinity for LCA, such as triantennary forms (Fig. 6B).

MALDI-TOF analysis of N-glycans released from Gb3/CD77 synthase produced in High Five cells showed the presence of oligomannose and hybrid structures (corroborating lectin blotting) (Fig. 7). Peaks corresponding to paucimannose N-glycans (main peak at *m/z* 1174.41, FucMan3GlcNAc2 followed by *m/z* 1028.36, Man3GlcNAc2) are predominant. Moreover, the spectrum shows weak signals derived from oligomannose structures (Man5GlcNAc2, Man6GlcNAc2, Man7GlcNAc2, Man8GlcNAc2, Man9GlcNAc2) and hybrid structures (*m/z* 1742.66, GalGlcNAc2FucMan3GlcNAc2) (Table S3). Contrary to SNA lectin blot analysis, no sialylated structures were detected, indicating that the weak SNA staining was non-specific (Fig. 6A, Table S3).

**Figure 7.**
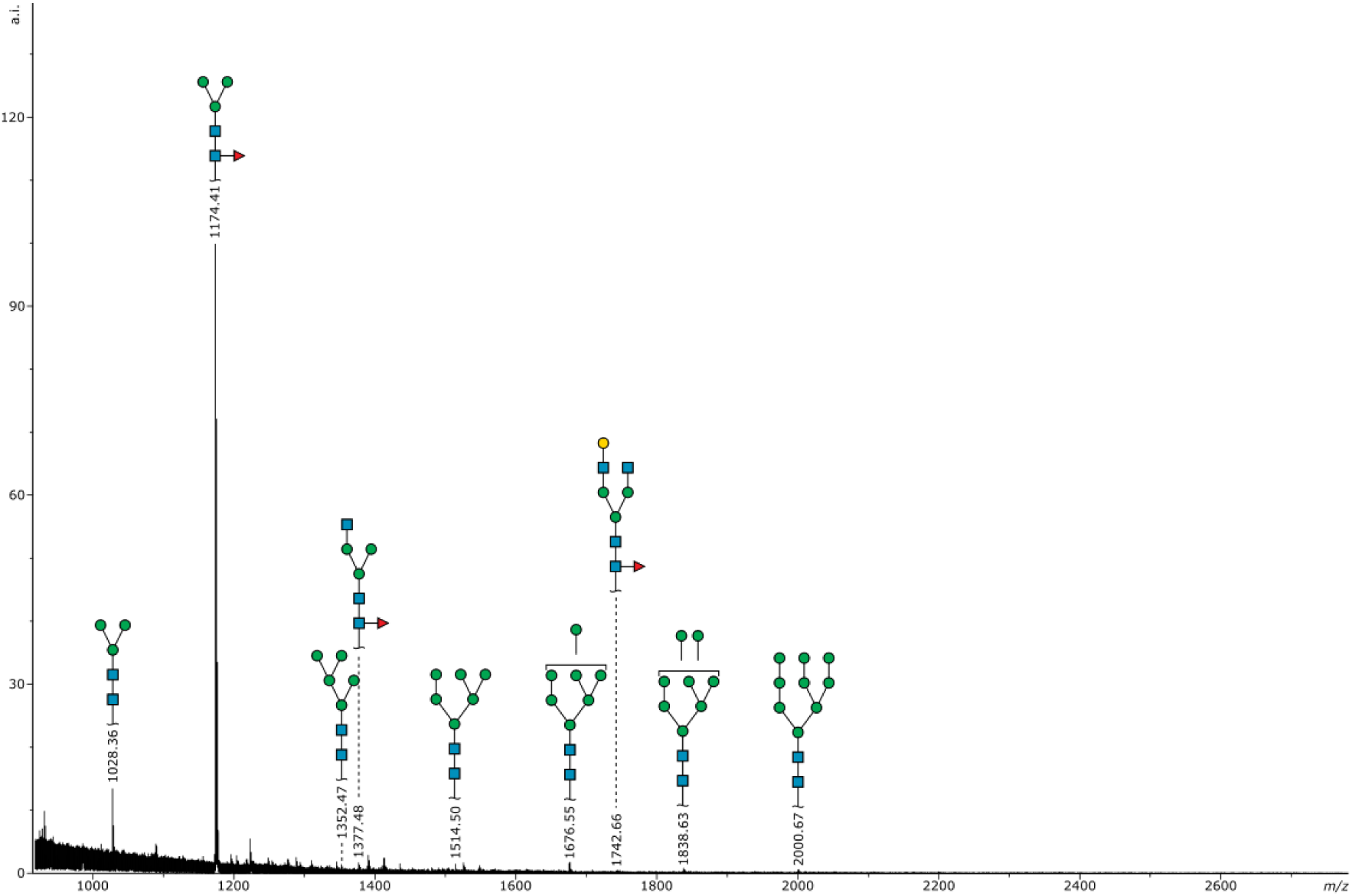
MALDI-TOF spectrum of 2-AA labeled N-glycans derived from the recombinant catalytic domain of human Gb3/CD77 synthase produced in High Five cells. All molecular ions are present in [M-H]^−^ form. The structures were annotated based on the putative composition and biosynthetic knowledge. Symbol representation is applied according to Symbol Nomenclature for Glycans (SNFG) [42].

MALDI-TOF spectrum obtained for N-glycans released from Gb3/CD77 synthase produced in Expi293F cells is rich in signals derived from complex, fucosylated structures (Fig. 8). Main peaks correspond to the triantennary monosialylated, difucosylated glycan (*m/z* 2382.86), biantennary difucosylated (*m/z* 2212.76) and triantennary, trifucosylated glycans (*m/z* 2237.82) (Table S4). Additionally, the spectrum presents signals derived from a variety of fucosylated bi-, tri- and tetraantennary structures, with up to 2 sialic acids, as well as high mannose structures (Table S4). In light of these results, we presume that weak staining with LCA lectin blot (Fig. 6B, Table S4) was due to the lower interaction of LCA with fucosylated triantennary glycans, which are prevalent among glycans released from the enzyme produced in Expi293F cells.

**Figure 8.**
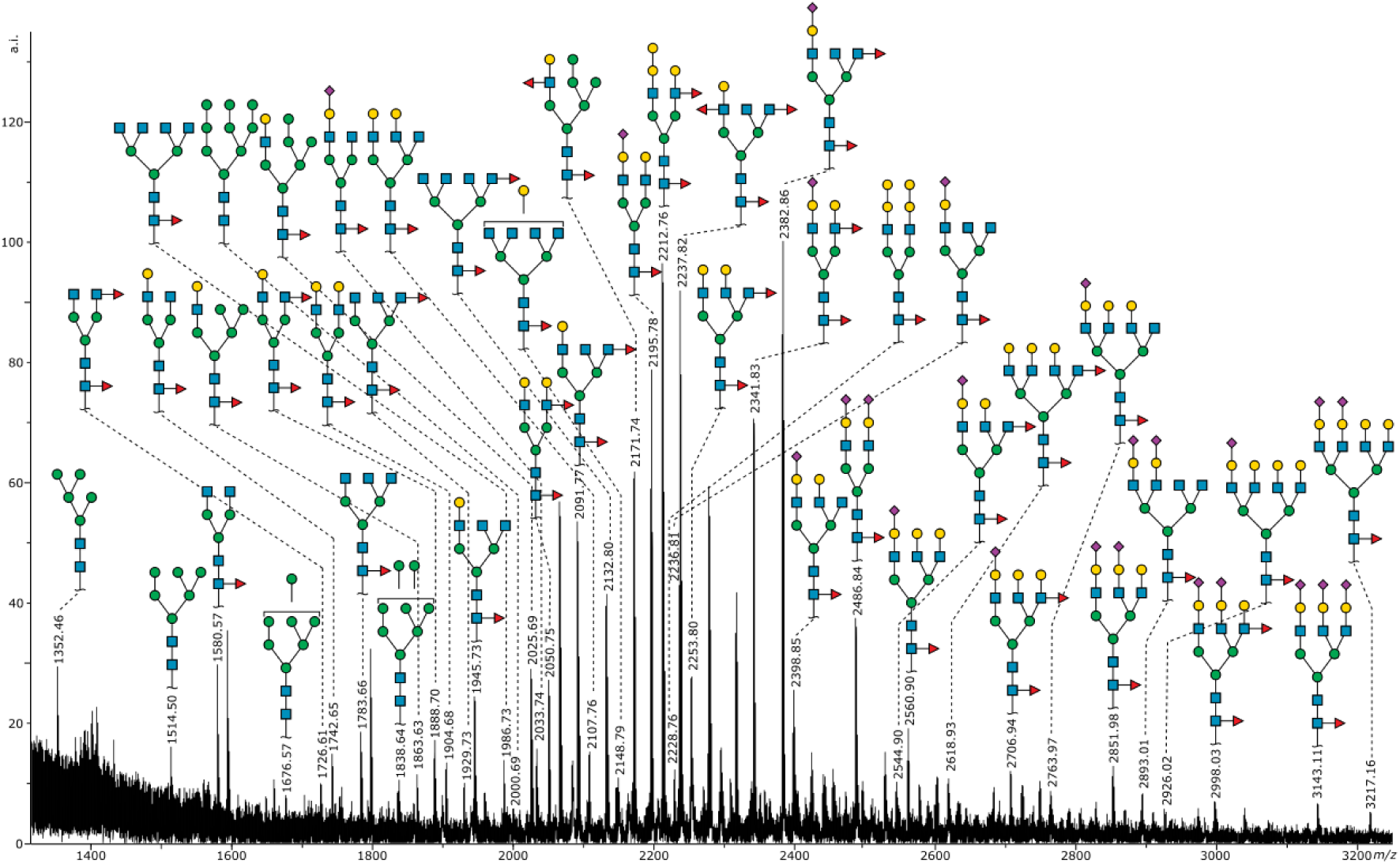
MALDI-TOF spectrum of 2-AA labeled N-glycans derived from the recombinant catalytic domain of human Gb3/CD77 synthase produced in Expi293F cells. All molecular ions are present in [M-H]^−^ form. The structures were annotated based on mass-based composition and biosynthetic knowledge. Symbol representation is applied according to Symbol Nomenclature for Glycans (SNFG) [42].

## Discussion

Recombinant glycosyltransferases have significant applications in various fields, such as in the production of biopharmaceuticals (therapeutic proteins, vaccine-related antigens, and monoclonal antibodies), applied biotechnology and biomedicine (for example, biomarkers development) [13–15]. In addition, recombinant GTs provide critical insights into the molecular determinants of enzyme specificity, kinetics, and substrate promiscuity, which are fundamental for understanding glycosylation-related biological processes [11,17,18].

Most GTs are membrane-associated proteins displaying high hydrophobicity, which makes their production in active form, combined with high purity and yield, outstandingly challenging. Therefore, a key consideration in recombinant GT production is the selection of an appropriate expression system, particularly taking into account that GTs often require PTMs, such as glycosylation, for correct folding and activity. Although prokaryotic hosts, such as *E. coli,* are widely used due to their simplicity and high yield of production, they frequently lack the machinery for essential PTMs, leading to the production of inactive enzymes, as demonstrated for MGAT3 [19] or MGAT4 [20]. Molecular chaperones may help proteins fold, such as in the case of α2,6-sialyltransferase ST6Gal-I, successfully expressed in *E. coli* strain pGro7/BL21, where the enzyme expression was driven by a cold-shock promoter and the GroEL/GroES chaperones by L-arabinose. [21]. However, eukaryote-based expression systems, such as invertebrate: insect cells *Spodoptera frugiperda* (Sf9), *T. ni* (High Five) [10,21–24], as well as vertebrate, such as baby hamster kidney cells (BHK21) [26], or human embryonic kidney cells (HEK293) may be more suitable for PTM-demanding proteins [8]. An alternative to animal cell culture expression systems is the Leishmania Expression System (LEXSY), which utilizes the protist *Leishmania tarentolae*, and was successfully used to produce several proteins [26–29]. LEXSY combines key advantages of mammalian systems (the ability to synthesize most PTMs, including complex N-glycans) with typical prokaryotic-related systems (such as culturing simplicity, low cost and high yield in protein production) [9].

In this study, we compared the expression of the soluble catalytic domain of human Gb3/CD77 synthase using two animal hosts: insect cells *T. ni* (High Five) and human-derived Expi293F cells. The highest yield of soluble human Gb3/CD77 synthase was obtained using High Five insect cells, while Expi293F cells exhibited lower production efficiency (Table 1, Fig. 9). The soluble human Gb3/CD77 synthase expressed in Expi293F cells appeared at approximately 50 kDa, whereas the enzyme from High Five cells migrated at around 40 kDa. This difference in apparent molecular weights is due to host-specific N-glycosylation patterns [31–33]. Using mass spectrometry and lectin blotting, we found that the enzyme from Expi293F cells carries complex-type N-glycans, including sialylated and multi-antennary structures, while the enzyme from High Five cells contains paucimannose and oligomannose glycans. Both systems yielded a catalytically active enzyme. In the case of the Expi293F-derived enzyme, activity was dependent on the localization of the 6xHis-tag; only the N-terminally tagged variant was active, while C-terminal tagging resulted in complete loss of catalytic function.

**Figure 9.**
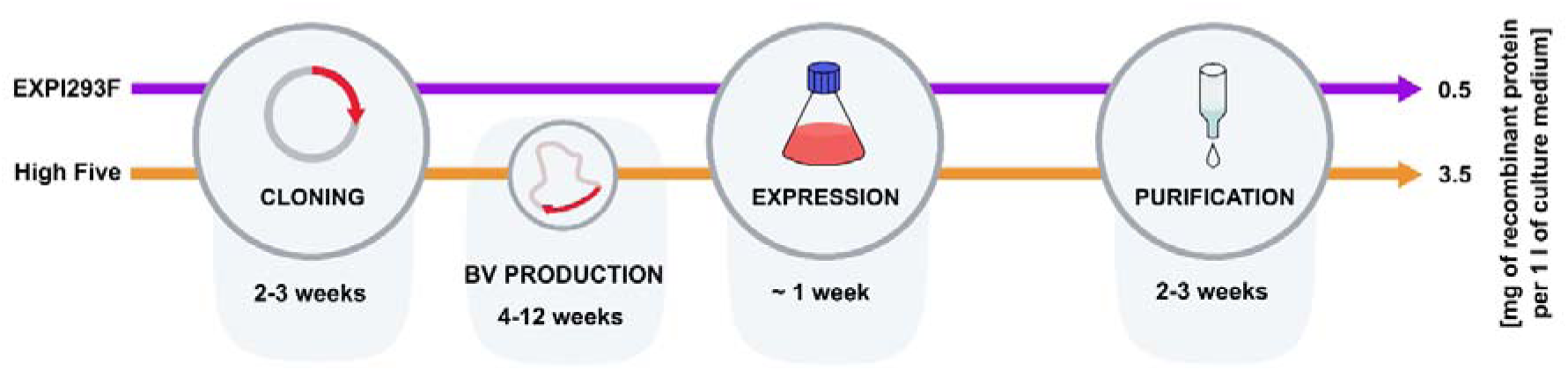
A workflow diagram presenting the time consumption and efficiency of the two expression hosts.

The lack of activity of the C-terminally tagged Expi293F-derived enzyme may be caused by a steric interference of the tag with the catalytic center. We hypothesize that positioning the tag at the N-terminus, which replaces the omitted transmembrane domain, results in less impact on protein folding and enzymatic function compared to C-terminal tagging. This strategy may thus be more appropriate for type II transmembrane enzymes, such as human Gb3/CD77 synthase. Recently, this principle was applied in the generation of an expression vector library encoding all known human glycan-modifying enzymes [32]. In this study, the catalytic domains of glycosyltransferase, glycosidases and sulfotransferases were produced in insect Sf9 and human HEK293 cells as soluble, GFP-fusion proteins. While additional tags were used for protein purification, the GFP domain enabled the quantitation of enzyme expression levels both inside the cells and when secreted into the media. The N-terminally tagged catalytic domain of Gb3/CD77 synthase, obtained in that study, showed high secretion efficiency in both insect and human expression systems; however, its activity was not evaluated. Instead, here we present the production of active, C-terminally tagged Gb3/CD77 synthase in High Five cells (Table 1 and Fig. 5A).

Human β1,4-galactosyltransferase 1 (B4GALT1), also a type II transmembrane protein, was produced in soluble form in the silkworm-based baculovirus expression system either N- or C-terminally tagged [34]. Interestingly, although the C-terminally tagged enzyme exhibited higher expression levels, its activity was slightly reduced compared to the N-terminally tagged variant. It suggests that tag positioning may have a more pronounced influence on GT activity in mammalian expression systems than in invertebrate systems, such as the silkworm-derived BEVS [32]. Moreover, elucidating the precise impact of tag orientation on enzymatic function will require detailed structural studies [35].

In both High Five and Expi293F cells, we obtained the glycosylated catalytic domain of human Gb3/CD77 synthase, with N-glycan profiles corresponding to the host-specific glycosylation machinery (paucimannose type dominating in High Five and complex in Expi293F). N-glycan structure is known to affect enzymatic activity, as demonstrated in human ST6Gal1, where the enzyme expressed in *Pichia pastoris* (producing short high-mannose glycans) was active, whereas the same enzyme expressed in *Saccharomyces cerevisiae* (producing long, branched glycans) was inactive [31]. Recently, human β1,3-galactosyltransferase 5 isoenzyme 1 (β3GalT5-1), which has three occupied N-glycosylation sites, was produced in soluble form in *E. coli, P. pastoris*, insect Sf9 and HEK293 cells [36]. Protein produced in *E. coli* and *P. pastoris* did not show enzymatic activity, presumably due to misfolding. Notably, enzymatically active β3GalT5-1 was obtained in *E. coli* only when co-expressed with molecular factors that promote disulfide bond formation. Conversely, β3GalT5-1 produced in insect Sf9 cells (carrying paucimannose glycans) and HEK293 cells (carrying complex glycans) demonstrated approximately ten times higher efficiency at producing glycans in comparison to the *E. coli*-derived enzyme [34].

Recently, we showed that glycosylation is fundamental for the secretion and activity of recombinant human Gb3/CD77 synthase [7,8]. For its enzymatic activity, an N-glycan at position N_203_ acts as a crucial “on” switch, while the N-glycan at the N_121_ site is an additional modulator [7]. We cannot exclude the influence of host-specific N-glycosylation pattern on recombinant Gb3/CD77 synthase folding, thereby affecting catalytic activity [8]. Perhaps the profound difference in glycosylation also underlies the discrepancy in the activity of C-tagged enzymes produced in High Five and Expi293F cells.

Both the High Five and Expi293F expression systems were able to produce catalytically active Gb3/CD77 synthase, but these hosts differed in yield, purity, glycosylation profile and dimerization capacity of the recombinant enzyme (Table 1). High Five cell cultures yielded a higher amount of soluble recombinant enzyme (approx. 3.5 mg/L) than Expi293F cultures (about 0.5 mg/L) (Table 1, Fig. 9). Only High Five-derived recombinant Gb3/CD77 synthase demonstrated the ability to form dimers, and enzyme produced in Expi293F cells was exclusively monomeric, suggesting that the expression host may influence the oligomeric state of the enzyme. High Five cells offer key advantages in cost-effectiveness, scalability and suitability in high-yield protein production. In contrast, Expi293F cells, although providing a more complex PTM repertoire, show limitations including lower expression yield, reduced purification efficiency and higher production costs [9]. These findings emphasize the importance of host selection in the production of recombinant GT and pave the way for further optimization strategies tailored to the intended application of the enzyme. A comparative summary of the expression outcomes is presented in Table 1 and Fig. 9.

Our data offer important learnings with broad implications for recombinant protein production in the context of gene therapy and the manufacture of biologics. The striking disparities in the functionality of tagged enzymes across cellular systems highlight the critical impact of the expression system on protein folding and biological activity. Our N-tagged enzymes derived from Expi293F cells exhibited robust activity, whereas their C-tagged counterparts were inactive. Intriguingly, both tags yielded active enzymes when expressed in insect cells. On the other hand, C-tagged glycosyltransferases were used for oligomerization studies in COS7 cells because N-tagged variants were prone to retention in the endoplasmic reticulum (ER), suggestive of misfolding [37]. However, the enzymatic activity of the C-terminally tagged variants was not evaluated in that study. These observations underscore three pivotal learnings: (1) expression systems differ markedly in their capacity to fold and secrete biologically active proteins; (2) cellular secretory pathways vary in efficiency, influencing protein trafficking; and (3) successful journey from ribosome to extracellular space through the ER and Golgi does not guarantee proper folding and/or functionality [37]. Incompatibility between target cells and the protein of interest has been implicated in suboptimal outcomes of gene therapy for hemophilia A, where precise protein folding and activity are paramount for therapeutic efficacy [38].

The recombinant catalytic domain of human Gb3/CD77 synthase obtained in this study was previously used for acceptor specificity evaluation [11], elucidating the role of N-glycosylation in the enzyme activity [8] and *ex vivo* glycosylation of human erythrocytes to modify their P1PK phenotype [39]. In future applications, the recombinant human Gb3/CD77 synthase holds promise biomanufacturing of anti-Stx agents, particularly via synthesis of the P1 glycotope (Galα1→4Galβ1→4GlcNAc-R)-bearing molecules aimed at treatment and prevention of Stxs-associated diseases [3,40]. To exploit the potential of these applications, further optimization of expression strategies is essential, particularly to ensure high yield, native-like PTMs, correct protein folding, and high purity of the final recombinant enzyme.

## Acknowledgments

The figures in the manuscript were created using BioRender.com.

## Data availability

Raw data, peak lists and identification results are available at GlycoPOST under ID: GPST000587 [41].

---

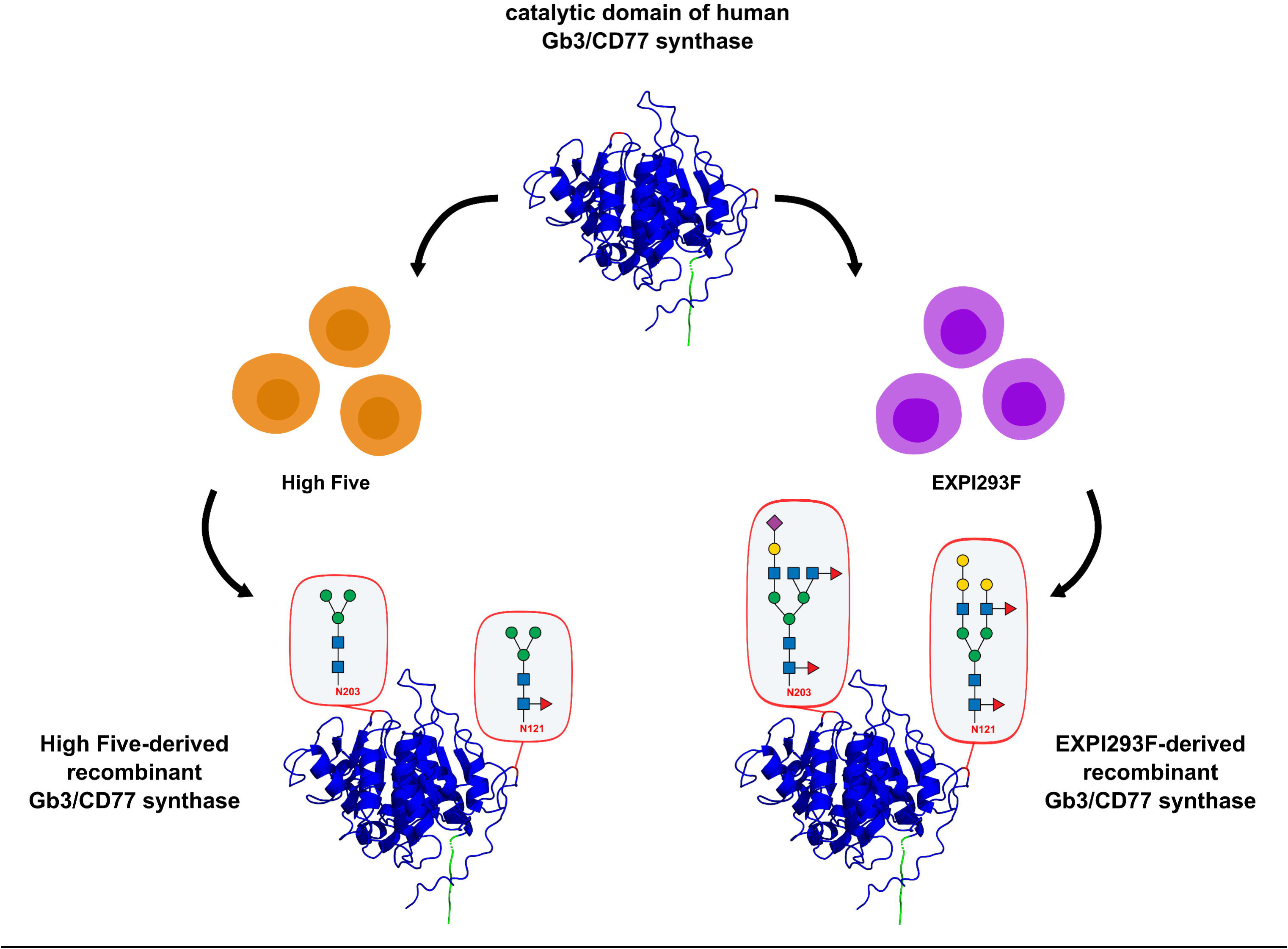

## Notes

### Competing Interest Statement

The authors have declared no competing interest.

### Summary of Updates

Abstract updated; Introduction, Results and Discussion sections revised; added new references; SI file updated.

