## Supplementary Material for "Expression of a human Gb3/CD77 synthase in insect and human cells: comparison of activity and glycosylation"

**Table S1**. List of primers used in this study. Restriction sites included in the primers used for cloning were underlined.

| **Name of primer** | **Sequence [5' → 3']** |
| --- | --- |
| **High Five** | |
| A4GTSeqsense | GGAGAGCCCAAGGAGAAAG |
| A4GTSeqanti | CAAGTACATTTTCATGGCCTC |
| A4GTBVsense | CATGGGAATTCCGGAGAGCCCAAGGAGAAAG |
| A4GTBVanti | AAAAGCGGCCGCCTAATGATGATGATGATGATGCAGATCCTCTTCTGAGATGAGTTTTTGTTCCAAGTACATTTTCATGGCCTC |
| **EXPI293F** | |
| A4GHEKNTAGsense | AAAAAAGCGGCCGCCCATCACCATCACCATCACTGGAGCCATCCTCAGTTTGAAAAGTCCGCTGGAGAGCCCAAGGAGAAAG |
| A4GHEKNTAGanti | AAAAAGAATTCTCACAAGTACATTTTCATGGCCTC |
| A4GHEKCTAGsense | AAAAAAGCGGCCGCCGGAGAGCCCAAGGAGAAAG |
| A4GHEKCTAGanti | AAAAAAGAATTCTCAGTGATGGTGATGGTGATGCTTTTCAAACTGAGGATGGCTCCAAGCGGACAAGTACATTTTCATGGCCTC |

**Table S2.** Lectin specificities and buffer compositions.

| **Lectin** | **Recognized carbohydrate structure** | **Buffer composition** | **Lectin manufacturer** |
| --- | --- | --- | --- |
| Concanavalin A, *Canavalia ensiformis* agglutinin (Con A) | Oligomannose- and hybrid-type N-glycans, and weakly biantennary complex-type N-glycans; trimannosidic N-glycan core with or without bisecting β1-->4-linked GlcNAc | TBS/0.05% Tween 20, pH 6.3, 1 mM CaCl_2_, 1 mM MnCl | Vector Laboratories, Inc., Newark, CA, USA |
| *Lens culinaris* agglutinin (LCA) | bi- and triantennary complex-type N-glycan with α1-6-linked core fucose | TBS/0.05% Tween 20, pH 7.4, 1 mM CaCl_2_, 1 mM MnCl | Vector Laboratories, Inc., Newark, CA, USA |
| *Sambucus nigra* agglutinin (SNA) | α2-->6-linked sialic acid, 6-O-sulfated LacNAc | TBS/0.05% Tween 20, pH 5.5 | Vector Laboratories, Inc., Newark, CA, USA |
| *Ricinus communis* I agglutinin (RCA I) | β-linked terminal Gal | TBS/0.05% Tween 20, pH 7.4 | Vector Laboratories, Inc., Newark, CA, USA |

**Table S3**. Compositions of N-glycans released from Gb3/CD77 synthase produced in High Five cells by MALDI-TOF-MS. H, hexose; N, *N*-acetylhexosamine; F, deoxyhexose (Fuc). The tallest peak is shown in bold.

| Structure | *m/z* (glycan + 2-AA) [M-H]^-^ | | Difference between the observed and theoretical *m/z* |
| --- | --- | --- | --- |
|  | Theoretical | Observed |  |
| H3N2 | 1028.36 | 1028.36 | Calibrant (0) |
| **H3N2F1** | **1174.41** | **1174.41** | **Calibrant (0)** |
| H5N2 | 1352.46 | 1352.47 | Calibrant (+0.01) |
| H3N3F1 | 1377.49 | 1377.48 | −0.01 |
| H6N2 | 1514.53 | 1514.50 | Calibrant (-0.03) |
| H7N2 | 1676.57 | 1676.55 | Calibrant (-0.02) |
| H4N4F1 | 1742.63 | 1742.66 | Calibrant (+0.03) |
| H8N2 | 1838.62 | 1838.63 | Calibrant (+0.01) |
| H9N2 | 2000.67 | 2000.67 | Calibrant (0) |

**Table S4**. Compositions of N-glycans released from Gb3/CD77 synthase produced in EXPI293F cells by MALDI-TOF-MS. H, hexose; N, *N*-acetylhexosamine; F, deoxyhexose (Fuc); S, sialic acid (NeuAc). Peaks with a relative intensity above 85% are shown in bold.

| Structure | *m/z* (glycan + 2-AA) [M-H]^-^ | | Difference between the observed and theoretical *m/z* |
| --- | --- | --- | --- |
|  | Theoretical | Observed |  |
| H5N2 | 1352.46 | 1352.46 | Calibrant (0) |
| H6N2 | 1514.49 | 1514.50 | Calibrant (+0.01) |
| H3N4F1 | 1580.57 | 1580.57 | 0 |
| H7N2 | 1676.57 | 1676.57 | Calibrant (0) |
| H3N4F2 | 1726.63 | 1726.61 | -0.02 |
| H4N4F1 | 1742.63 | 1742.65 | Calibrant (+0.02) |
| H3N5F1 | 1783.65 | 1783.66 | +0.01 |
| H8N2 | 1838.62 | 1838.64 | Calibrant (+0.02) |
| H6N3F1 | 1863.65 | 1863.63 | -0.02 |
| H4N4F2 | 1888.68 | 1888.70 | +0.02 |
| H5N4F1 | 1904.68 | 1904.68 | Calibrant (0) |
| H3N5F2 | 1929.71 | 1929.73 | +0.02 |
| H4N5F1 | 1945.71 | 1945.73 | +0.02 |
| H3N6F1 | 1986.73 | 1986.73 | 0 |
| H9N2 | 2000.67 | 2000.69 | Calibrant (+0.02) |
| H7N3F1 | 2025.70 | 2025.69 | -0.01 |
| H4N4F1S1 | 2033.72 | 2033.74 | +0.02 |
| H5N4F2 | 2050.74 | 2050.75 | +0.01 |
| H4N5F2 | 2091.76 | 2091.77 | +0.01 |
| H5N5F1 | 2107.76 | 2107.76 | 0 |
| H3N6F2 | 2132.79 | 2132.80 | +0.01 |
| H4N6F1 | 2148.79 | 2148.79 | 0 |
| H7N3F2 | 2171.76 | 2171.74 | -0.02 |
| H5N4F1S1 | 2195.77 | 2195.78 | +0.01 |
| **H6N4F2** | **2212.79** | **2212.76** | **-0.03** |
| H7N4F1 | 2228.78 | 2228.76 | -0.02 |
| H4N5F1S1 | 2236.80 | 2236.81 | +0.01 |
| **H4N5F3** | **2237.82** | **2237.82** | **0** |
| H5N5F2 | 2253.82 | 2253.80 | -0.02 |
| H5N4F2S1 | 2341.83 | 2341.83 | 0 |
| **H4N5F2S1** | **2382.86** | **2382.86** | **0** |
| H5N5F1S1 | 2398.86 | 2398.85 | -0.01 |
| H5N4F1S2 | 2486.87 | 2486.84 | Calibrant (-0.03) |
| H5N5F2S1 | 2544.91 | 2544.90 | -0.01 |
| H6N5F1S1 | 2560.91 | 2560.90 | -0.01 |
| H6N6F2 | 2618.95 | 2618.93 | -0.02 |
| H6N5F2S1 | 2706.96 | 2706.94 | -0.02 |
| H6N6F1S1 | 2763.99 | 2763.97 | -0.02 |
| H6N5F1S2 | 2852.00 | 2851.98 | -0.02 |
| H5N6F1S2 | 2893.03 | 2893.01 | -0.02 |
| H7N6F1S1 | 2926.04 | 2926.02 | -0.02 |
| H6N5F2S2 | 2998.03 | 2998.03 | 0 |
| H6N5F1S3 | 3143.10 | 3143.11 | Calibrant (+0.01) |
| H7N6F1S2 | 3217.13 | 3217.16 | +0.01 |
